# Apolipoprotein E2 Promotes Melanoma Growth, Metastasis, and Protein Synthesis via the LRP1 Receptor

**DOI:** 10.1101/2022.10.03.510632

**Authors:** Nneoma Adaku, Benjamin N. Ostendorf, Sohail F. Tavazoie

## Abstract

The secreted lipid transporter apolipoprotein E (APOE) plays important roles in atherosclerosis and Alzheimer’s disease and has been implicated as a suppressor of melanoma progression. *APOE* germline genotype predicts human melanoma outcomes, as *APOE4* and *APOE2* allele carriers exhibit increased versus reduced melanoma survival, respectively, relative to *APOE3* homozygotes. While the APOE4 variant was recently shown to suppress melanoma progression by enhancing anti-tumor immunity, the melanoma cell-intrinsic effects of APOE variants on cancer progression remain poorly characterized. By using a genetically engineered mouse model, we show that human germline *APOE* genetic variants differentially modulate melanoma growth and metastasis in a melanoma LRP1 receptor-dependent manner. We identify protein synthesis as a tumor cell-intrinsic process differentially modulated by APOE variants, with APOE2 surprisingly promoting translation via LRP1. Our findings reveal a gain-of-function role for the APOE2 variant in melanoma progression, raising important implications for other diseases impacted by *APOE* genetics.

**Significance:** Our work reveals that germline *APOE* variants differentially impact genetically initiated melanoma progression, with APOE2 acting as a promoter of tumor growth, metastasis, and a cell-intrinsic process—protein synthesis. These findings may aid in predicting patient outcomes and may partly explain the protective effect of APOE2 in Alzheimer’s disease.

## Introduction

Germline genetic variants are well-established regulators of cancer development. They underlie hereditary cancer predisposition syndromes, which account for 5-10% of all malignancies (1), and an estimated 33% of all cancer is thought to be heritable (2). In contrast, the impact of germline genetics on the progression of cancers once they are established is poorly understood. We previously identified genetic variation in the secreted glycoprotein apolipoprotein E (APOE) as a significant modulator of survival after melanoma development (3). Humans have three prevalent *APOE* alleles, termed *APOE2, APOE3*, and *APOE4*, that differ by variation at two amino acid residues. In contrast to Alzheimer’s disease, where *APOE4* is the single greatest monogenetic risk factor for disease onset and *APOE2* is protective, melanoma patients who carry an *APOE4* allele exhibit improved survival whereas *APOE2* carriers experience poorer outcomes in multiple large cohorts. The variant-dependent effects of APOE on melanoma progression are partly governed by differences in anti-tumor immunity, as stromal expression of APOE4 confers increased immune effector responses relative to APOE2. Moreover, APOE4 has been shown to suppress tumor angiogenesis and endothelial recruitment by cancer cells (3,4). In addition to tumor cell-extrinsic effects on the immune and vascular compartments, prior work also identified a cell-intrinsic effect of APOE in suppressing of melanoma cell invasiveness (4). This cell-intrinsic effect was also variant dependent, with APOE4 most potently suppressing invasion relative to APOE2 (3). Overall, the effects of APOE4 including enhanced anti-tumor immunity, reduced angiogenesis, and reduced invasiveness collectively contribute to its melanoma-suppressive effects relative to APOE2. These past studies revealed an APOE4>APOE3>APOE2 pattern of melanoma tumor suppression. While this past work led to the identification of multiple cellular and organismal cancer progression phenotypes that are differentially associated with APOE variants, the impact of APOE variants on intracellular processes that could contribute to cancer growth and metastasis outcomes has remained uncharacterized.

Previous studies investigating the role of APOE in melanoma progression have primarily utilized transplantable models, in which established cancer cells are injected directly into mice (3-5). Herein, we crossed the *Braf*^*V600E*^/*Pten*^-/-^ conditional melanoma model (6) with mice in which the murine *Apoe* locus had been replaced with one of the three human *APOE* genes—thus generating an allelic series of genetically engineered mouse models (GEMMs) of melanoma harboring human *APOE2, APOE3*, or *APOE4*. These allelic strains vary from one another by just one or two amino acids. In contrast to transplantable models, GEMMs recapitulate all steps of the cancer progression cascade from tumor initiation through to metastatic colonization, and the genetics of the tumor match that of the host. We therefore reasoned that cancer cell exposure to allele-concordant host and tumoral APOE for the entire metastatic trajectory, as it occurs in patients, may enhance phenotypic expression of the variants’ cell-intrinsic effects. We found that genetically initiated melanoma growth and metastasis were regulated by *APOE* genotype in an *APOE2*>*E3*>*E4* manner. Additionally, we identified mRNA translation as a melanoma cell-intrinsic process that is modulated by APOE, with APOE2 acting as a promoter of protein synthesis. These effects were dependent on melanoma cell expression of low-density lipoprotein receptor-related protein 1 (LRP1), a receptor for APOE. In the Alzheimer’s field, it has long been debated whether *APOE2* is solely a loss-of-function allele with respect to its effects on neurodegeneration or whether it harbors gain-of-function properties (7). Our findings reveal clear gain-of-function effects for APOE2 in both protein translation and the promotion of melanoma progression. Through systematic interrogation of the effects of germline genetic variation of the *APOE* gene in mouse and patient tumors, our work uncovers a novel role for APOE in the regulation of protein synthesis via the LRP1 receptor and provides definitive evidence for hereditary regulation of cancer progression and metastasis by common human germline genetic variants.

## Results

### Human APOE variants differentially modulate melanoma progression in a genetically engineered mouse model

To model the impact of human APOE variants on all stages of melanoma progression, we crossed the well-established *Braf*^*V600E/+*^*;Pten*^*−/ −*^*;Tyr::CreER* (BPC) GEMM with APOE-targeted replacement (knock-in) mice, in which the endogenous murine *Apoe* locus has been replaced with one of the three human *APOE* genes (8-10) **(Fig. 1A)**. The BPC model enables tamoxifen-inducible deletion of the *Pten* tumor suppressor and activation of the *Braf*^*V600E*^ oncogene specifically in melanocytes, resulting in melanoma formation in 3-4 weeks with 100% penetrance and recapitulation of the entirety of the metastatic cascade (6). 4-hydroxytamoxifen (4-OHT) was applied topically to the lower backs of BPC*;APOE2* (BPC/APOE2), BPC*;APOE3* (BPC/APOE3), and BPC*;APOE4* (BPC/APOE4) mice. Melanoma onset occurred with shortest latency in BPC/APOE2 mice, followed by BPC/APOE3 mice and then BPC/APOE4 mice **(Figs. 1B, 1C)**. Mouse survival followed a similar pattern, with BPC/APOE2 mice having the shortest median survival at 42.5 days, BPC/APOE3 mice intermediate at 53.5 days, and BPC/APOE4 mice having the longest survival at 59.5 days **(Fig. 1D)**. We thus focused mainly on the *APOE2* and *APOE4* genotypes for the remainder of the study, as they produced the most divergent tumor phenotypes in the GEMM. In an independent cohort of mice whose tumor growth rate was tracked, melanomas grew significantly faster **(Fig. 1E)** and were significantly larger at the day 49 endpoint **(Fig. 1F)** in BPC/APOE2 mice relative to BPC/APOE4 mice—consistent with the aforementioned findings.

**Figure 1.**
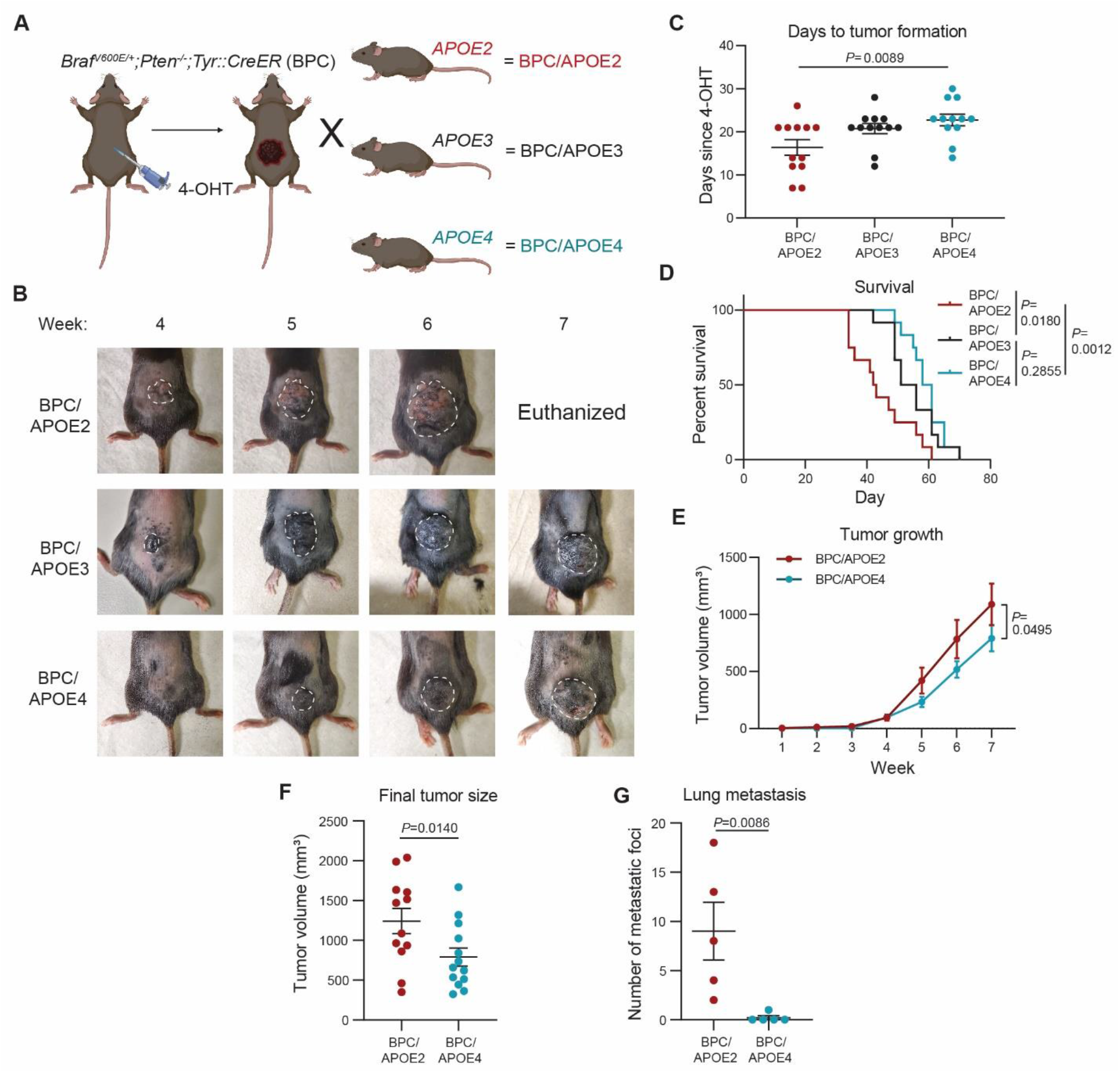
Common human APOE variants differentially impact melanoma growth and metastasis in a genetically engineered mouse model. **A**, Schematic depicting generation of, and tumor induction in, the *Braf* ^*V600E/+*^*;Pten*^*-/-*^*;Tyr::CreER*;*APOE2* (BPC/APOE2), ;*APOE3* (BPC/APOE3), and ;*APOE4* (BPC/APOE4) mouse models. **B**, Representative images of tumor growth in BPC/APOE2, BPC/APOE3, and BPC/APOE4 mice 4 to 7 weeks after topical administration of 4-OHT. **C**, Number of days after topical 4-OHT administration until tumors were palpated and visualized in BPC/APOE2, BPC/APOE3, and BPC/APOE4 mice (n=12 per group). One-way ANOVA. **D**, Kaplan-Meier survival curves of BPC/APOE2, BPC/APOE3, and BPC/APOE4 mice after topical 4-OHT administration (n=12 per group). Log-rank test. **E**, Tumor growth curve of BPC/APOE2 (n=12) and BPC/APOE4 (n=13) mice after topical 4-OHT administration. Two-way ANOVA. **F**, Final tumor volumes of BPC/APOE2 (n=12) and BPC/APOE4 (n=13) mice from **E** at the experimental endpoint of 49 days after topical 4-OHT administration. Unpaired t-test. **G**, Quantification of lung metastatic foci in BPC/APOE2 (n=5) and BPC/APOE4 (n=5) mice after neonatal tumor induction. Unpaired t-test.

Because APOE is a potent suppressor of melanoma metastasis (4), we next evaluated whether genetic variation in *APOE* could modulate metastatic capacity in the BPC GEMM. Metastatic burden in the BPC model can be determined by quantifying pigmented foci on the surface of the lung after neonatal administration of 4-OHT (11). Accordingly, BPC/APOE2 and BPC/APOE4 neonates received 4-OHT topically and were euthanized after weaning age. Pigmented foci were visible on the lungs of mice, and BPC/APOE2 mice exhibited substantially more lung metastatic burden compared to BPC/APOE4 mice **(Fig. 1G)**. These results in the APOE variant GEMM revealed a potent impact of hereditary genetics on tumor growth and metastatic colony formation in an autochthonous model of melanoma progression.

### Genetically engineered mouse model reveals protein translation upregulation in *APOE2* tumors

We next utilized the *APOE* allelic GEMM series as a tool to search for cellular processes that might be altered in an APOE variant-dependent manner and that could influence cancer progression. To this end, we performed bulk RNA-Seq of time-matched BPC/APOE2 and BPC/APOE4 tumors. Gene set enrichment analysis (GSEA) (12) revealed that translation was the most upregulated pathway in BPC/APOE2 tumors relative to BPC/APOE4 tumors **(Fig. 2A)**. The additional protein synthesis-related pathways “Ribosomal RNA processing” and “Eukaryotic translation elongation” were also among the top ten upregulated pathways. BPC/APOE2 tumors displayed significant upregulation in all major steps of translation present in the Reactome (13) gene set relative to BPC/APOE4 tumors **(Fig. 2B)**.

**Figure 2.**
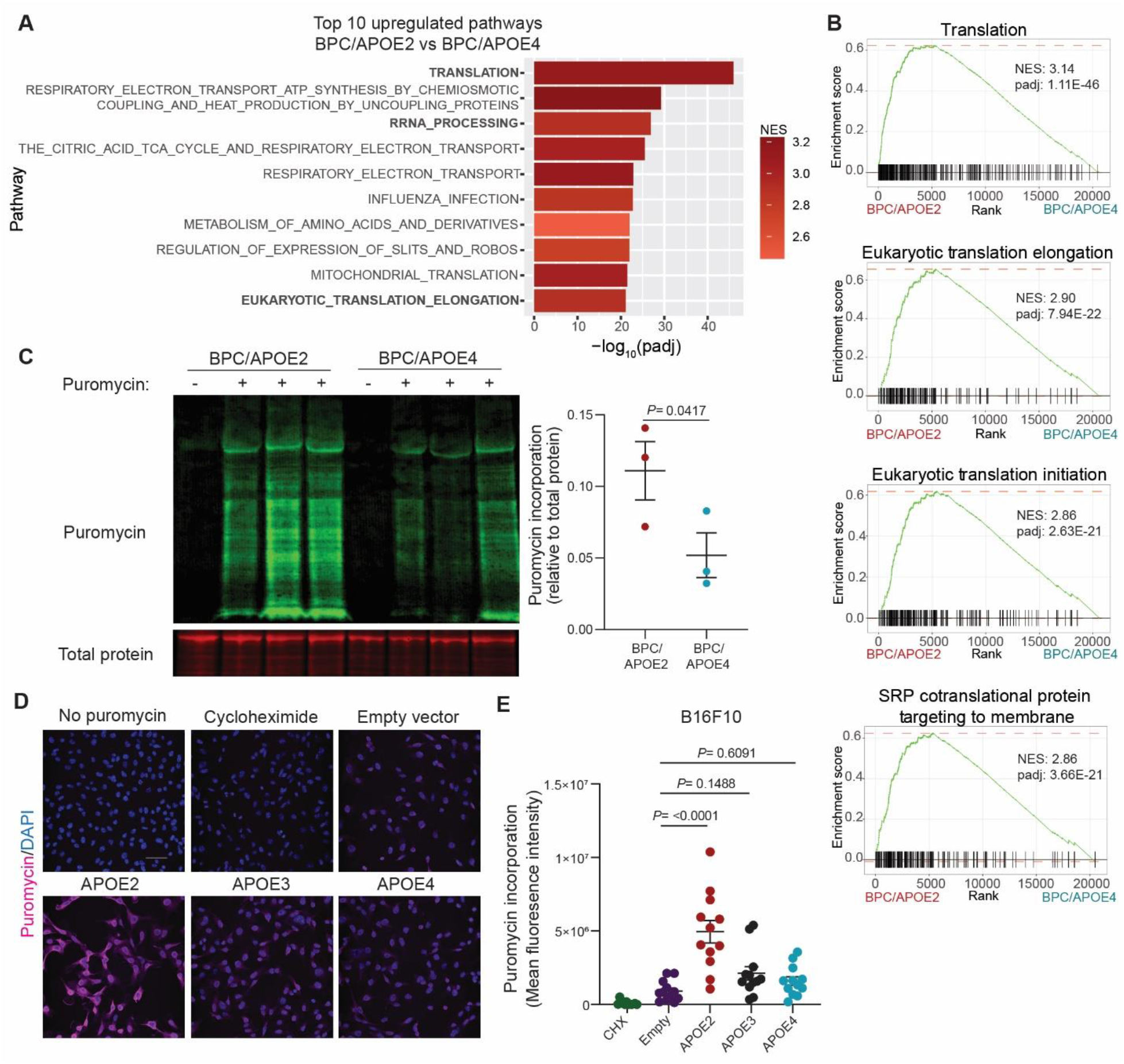
Common *APOE* germline genetic variants modulate protein synthesis in melanoma. **A**, Top ten pathways upregulated in melanomas of BPC/APOE2 mice relative to BPC/APOE4 mice as determined by GSEA and ranked by adjusted p-value (n=4 per group; NES, normalized enrichment score; padj, adjusted p-value). **B**, Enrichment plots of translation-related pathways within the Reactome gene set. **C**, Western blot of puromycin incorporation into BPC/APOE2 and BPC/APOE4 tumors 35 days after 4-OHT administration (n=3 per group). Non-puromycin-pulsed mice were included as an antibody control. Unpaired t-test. **D**, Representative immunofluorescence images of puromycin incorporation into B16F10-sh*Apoe* pBabe Empty, APOE3, APOE2, and APOE4 cells. Non-puromycin-pulsed and cycloheximide (CHX)-treated cells were included as negative and positive controls, respectively (scale bar = 50µM). **E**, Quantification of mean fluorescence intensity from SUnSET assay in **D**. One-way ANOVA (n=3 independent experiments).

Translational control plays a crucial role in all steps of cancer progression, and most oncogenic signaling pathways converge to enhance the translational capacity of tumor cells (14-16). We therefore sought to experimentally validate whether there were differences in translational efficiency between *APOE2* and *APOE4* melanomas. We utilized the surface sensing of translation (SUnSET) assay, a well-established method for measuring protein synthesis (17). In this assay puromycin can be administered to live mice, which then incorporates into nascent polypeptide chains synthesized in mouse tissues (18). Puromycin incorporation can then be quantified with an anti-puromycin antibody, providing a readout of global cellular translation. To control for tumor size differences, BPC/APOE2 and BPC/APOE4 mice were injected with puromycin 35 days after 4-OHT administration, an early time point at which a significant difference in tumor volumes was not yet detectable (Supplementary Fig. S1A). Consistent with our RNA-Seq results, BPC/APOE2 tumors exhibited significantly higher puromycin incorporation than BPC/APOE4 tumors, indicative of either a slower translation rate in the *APOE4* background or enhanced translation rate in the *APOE2* background **(Fig. 2C)**. To further control for differences in tumor proliferation and the presence of stromal cells, we performed the SUnSET assay *in vitro* with melanoma cells stably overexpressing APOE2, APOE3, APOE4, or an empty control vector (Supplementary Fig. 1B). Strikingly, there was no difference in puromycin incorporation between control, APOE3, and APOE4 cells, whereas APOE2 cells exhibited substantially increased puromycin signal **(Figs. 2D and 2E)**. These findings are consistent with a model whereby APOE2 promotes translation rather than APOE4 inhibiting translation.

We next sought orthogonal support of a protein translation role for APOE2 and also investigated whether the APOE-dependent effects on translation are present in metastatic disease. To do this, we performed tail vein injections of syngeneic, GFP-expressing B16F10 melanoma cells depleted of murine APOE into *APOE2* or *APOE3* knock-in mice. APOE3 served as an ideal isogenic control for determining whether APOE2 mediates translational enhancement because it exhibited an intermediate phenotype in our GEMM studies and is the most common APOE variant in the human population (7). The injections were followed by bulk RNA-Seq of metastatic melanoma cells that were isolated from lungs by flow cytometry (Supplementary Fig. 1C). *APOE2* mice had a significantly higher fraction of GFP+ tumor cells within their dissociated lungs relative to *APOE3* mice, indicative of higher metastatic burden (Supplementary Fig. 1D). Importantly, translation-related processes again dominated the top ten upregulated pathways in *APOE2* metastases relative to *APOE3* (Supplementary Fig. 1E). These results indicate that the APOE variants differentially impact melanoma protein synthesis, with APOE2 stimulating translation in both localized and metastatic melanoma.

### LRP1 mediates cell-intrinsic effects of APOE variants on melanoma progression

We previously found that the APOE receptor LRP1 mediates the cell-intrinsic effects of APOE on melanoma cells (4). We thus investigated whether LRP1 is necessary for the differential impact of APOE variants on melanoma progression. After tail vein injection, control B16F10 melanoma cells depleted of murine APOE metastasized to the lung more efficiently in *APOE2* knock-in mice relative to *APOE4* mice **(Fig. 3A)**. However, after CRISPR/Cas9-mediated deletion of melanoma *Lrp1* via two independent guide RNAs (Supplementary Fig. 2A), the difference in metastatic burden between *APOE2* and *APOE4* mice was abrogated **(Fig. 3B**, Supplementary Fig. 2B). These findings reveal that the effects of APOE variants on melanoma metastatic capacity are mediated via the APOE receptor LRP1.

**Figure 3.**
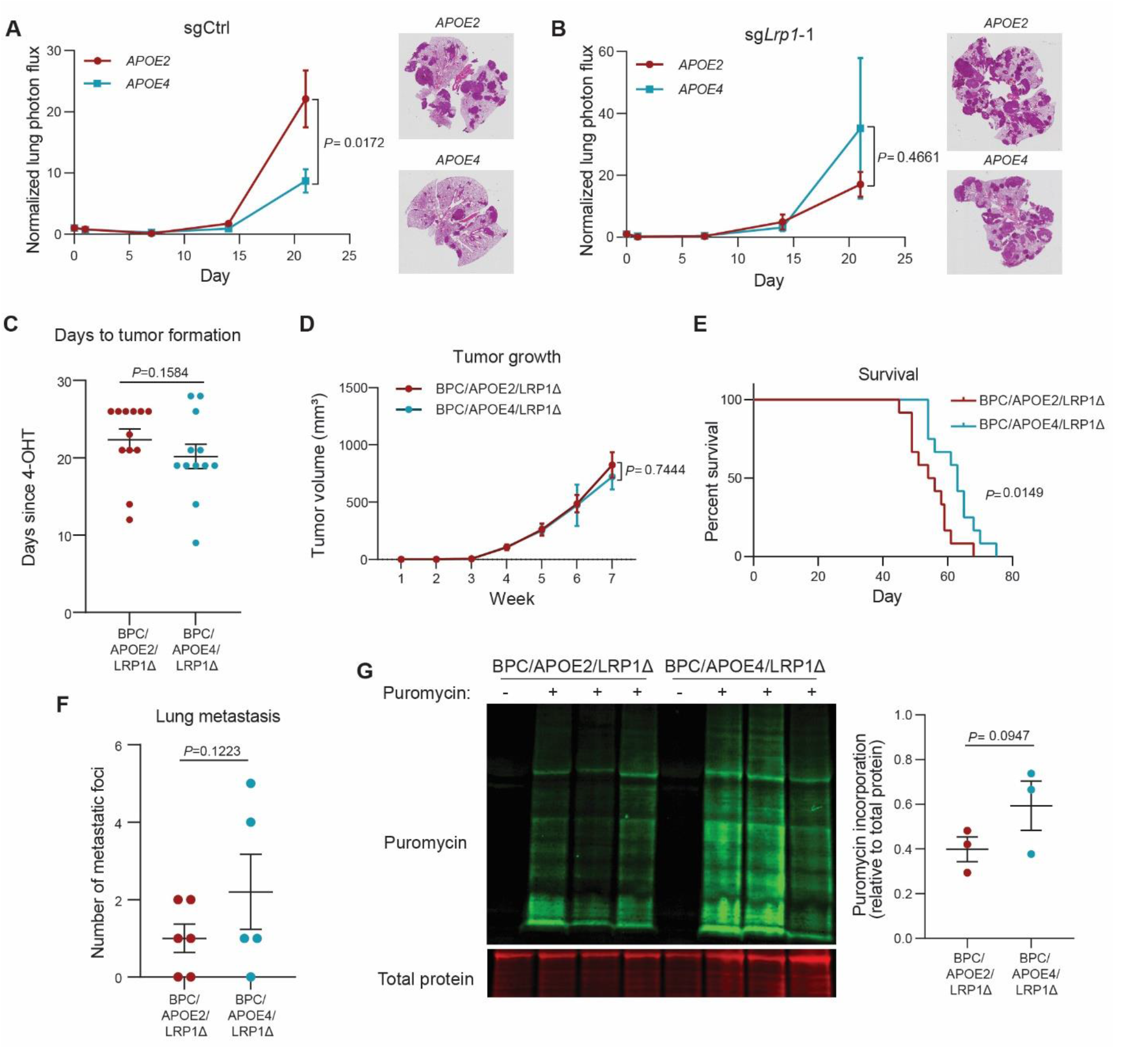
Differential effects of APOE variants on melanoma progression and protein translation are abrogated upon tumoral *Lrp1* genetic deletion. **A, B**, Quantification of lung metastatic progression via bioluminescence imaging of B16F10-TR-sh*Apoe* sgCtrl **(A)** or sg*Lrp1*-1 **(B)** cells injected via lateral tail vein into *APOE2* and *APOE4* mice. Representative images of H&E-stained lungs taken from mice at the day 21 endpoint (n = 9-10 mice per group; representative of two independent experiments; two-way ANOVA). **C**, Number of days after topical 4-OHT administration until tumors were palpated and visualized in BPC/APOE2/LRP1Δ and BPC/APOE4/LRP1Δ mice (n=12 per group). Unpaired t-test. **D**, Melanoma tumor growth curves of BPC/APOE2/LRP1Δ (n=12) and BPC/APOE4/LRP1Δ (n=10) mice after topical 4-OHT administration. Two-way ANOVA. **E**, Kaplan-Meier survival curves of BPC/APOE2/LRP1Δ and BPC/APOE4/LRP1Δ mice after topical 4-OHT administration (n=12 per group). Log-rank test. **F**, Quantification of lung metastatic foci in BPC/APOE2/LRP1Δ (n=6) and BPC/APOE4/LRP1Δ (n=5) mice after neonatal tumor induction. Unpaired t-test. **G**, Western blot of puromycin incorporation into BPC/APOE2/LRP1Δ and BPC/APOE4/LRP1Δ tumors 35 days after 4-OHT administration (n=3 per group). Non-puromycin-pulsed mice were included as an antibody control. Unpaired t-test.

Recognizing the limitations of transplantable models as described earlier, we crossed BPC/APOE2 and BPC/APOE4 mice with *Lrp1*^*flox/flox*^ mice, thus enabling deletion of *Lrp1* specifically in the melanocytes that go on to form melanomas after 4-OHT administration.

Immunofluorescence staining of BPC/APOE2*;Lrp1*^*flox/flox*^ (BPC/APOE2/LRP1Δ) and BPC/APOE4*;Lrp1*^*flox/flox*^ (BPC/APOE4/LRP1Δ) tumors showed substantial loss of LRP1 signal compared to *Lrp1* wild-type tumors, confirming successful Cre-mediated deletion (Supplementary Fig. S3A). We again monitored tumor growth in this model after 4-OHT administration to back skin of mice. In contrast to the differential effects of APOE variants observed in *Lrp1* wild-type mice, there was no significant difference in tumor latency **(Fig. 3C)**, tumor growth rate **(Fig. 3D)**, or tumor volume at the experimental endpoint (Supplementary Fig. S3B) between BPC/APOE2/LRP1Δ and BPC/APOE4/LRP1Δ mice. In a separate survival experiment, there was a significant difference with BPC/APOE2/LRP1Δ mice having a median survival of 55 days compared to 63 days in BPC/APOE4/LRP1Δ mice **(Fig. 3E)**. However, this difference was diminished compared to *Lrp1* wild-type BPC/APOE2 and BPC/APOE4 mice **(Fig. 1D)**. This result is consistent with our prior findings revealing that the *APOE4* background confers more potent anti-tumor immunity and repressed angiogenesis (3), thus providing BPC/APOE4/LRP1Δ mice a survival advantage over BPC/APOE2/LRP1Δ mice at later primary tumor stages. There was also no significant difference in distant lung metastasis between BPC/APOE2/LRP1Δ and BPC/APOE4/LRP1Δ mice after neonatal 4-OHT administration **(Fig. 3F)**, consistent with cell-intrinsic effects of APOE dominating at the metastatic site. These results indicate that *Lrp1* deletion significantly abrogates the impact of APOE variants on melanoma progression, with differences only emerging when the cell-extrinsic effects of APOE become impactful at later primary tumor stages in the genetic model.

Having determined that LRP1 mediates the cell-intrinsic effects of the APOE variants, we next investigated whether the effects of APOE variants on mRNA translation are LRP1-dependent. The SUnSET assay was performed with day 35 BPC/APOE2/LRP1Δ and BPC/APOE4/LRP1Δ tumors and showed equalization of puromycin incorporation between the genotypes upon *Lrp1* deletion **(Fig. 3G)**. These findings reveal that the APOE receptor LRP1 is a required mediator of the cell-intrinsic effects of APOE variants on melanoma progression at the early primary tumor and metastasis stages. Moreover, LRP1 is required for the enhanced protein synthesis effect mediated by APOE2 in melanoma.

### *APOE* genetics impact translation regulation in human melanomas

We next sought to determine whether the APOE variant-dependent translation phenotype could also be observed in human melanomas. Given the low allele frequency of *APOE2* (7), which limits statistical power, we turned to public datasets. The Cancer Genome Atlas skin cutaneous melanoma (TCGA-SKCM) cohort contains over 400 patients with both RNA sequencing and whole exome sequencing data, enabling differential gene expression analysis of human tumors based on the germline *APOE* genotype (19) **(Fig. 4A)**. The TCGA-SKCM dataset was previously used to determine that *APOE4* carriers have a survival advantage relative to *APOE3* patients while *APOE2* carriers experience worse survival outcomes (3). Consistent with our findings in BPC/APOE2 and BPC/APOE4 mice, primary tumors from *APOE2* carrier patients exhibited upregulation of translation pathways relative to tumors from *APOE4* carriers **(Fig. 4B)**. Also consistent with the notion that APOE2 is an active promoter of protein synthesis, translation pathways were also the top upregulated processes in *APOE2* carrier primary tumors compared to *APOE3* homozygotes (Supplementary Figs. 4A and 4B). Protein translation upregulation in the *APOE2* background relative to *APOE4* was also maintained in metastatic tumors, suggesting that APOE2 promotes protein synthesis in the melanomas of *APOE2* carriers throughout the metastatic cascade **(Fig. 4C)**. Taken together with our findings from mouse modeling, we propose that APOE2-mediated enhancement of translation—a key driver of tumor growth and metastasis—supports melanoma growth and metastasis in *APOE2* carriers, consistent with their poor survival compared to both *APOE3* homozygotes and *APOE4* carriers.

**Figure 4.**
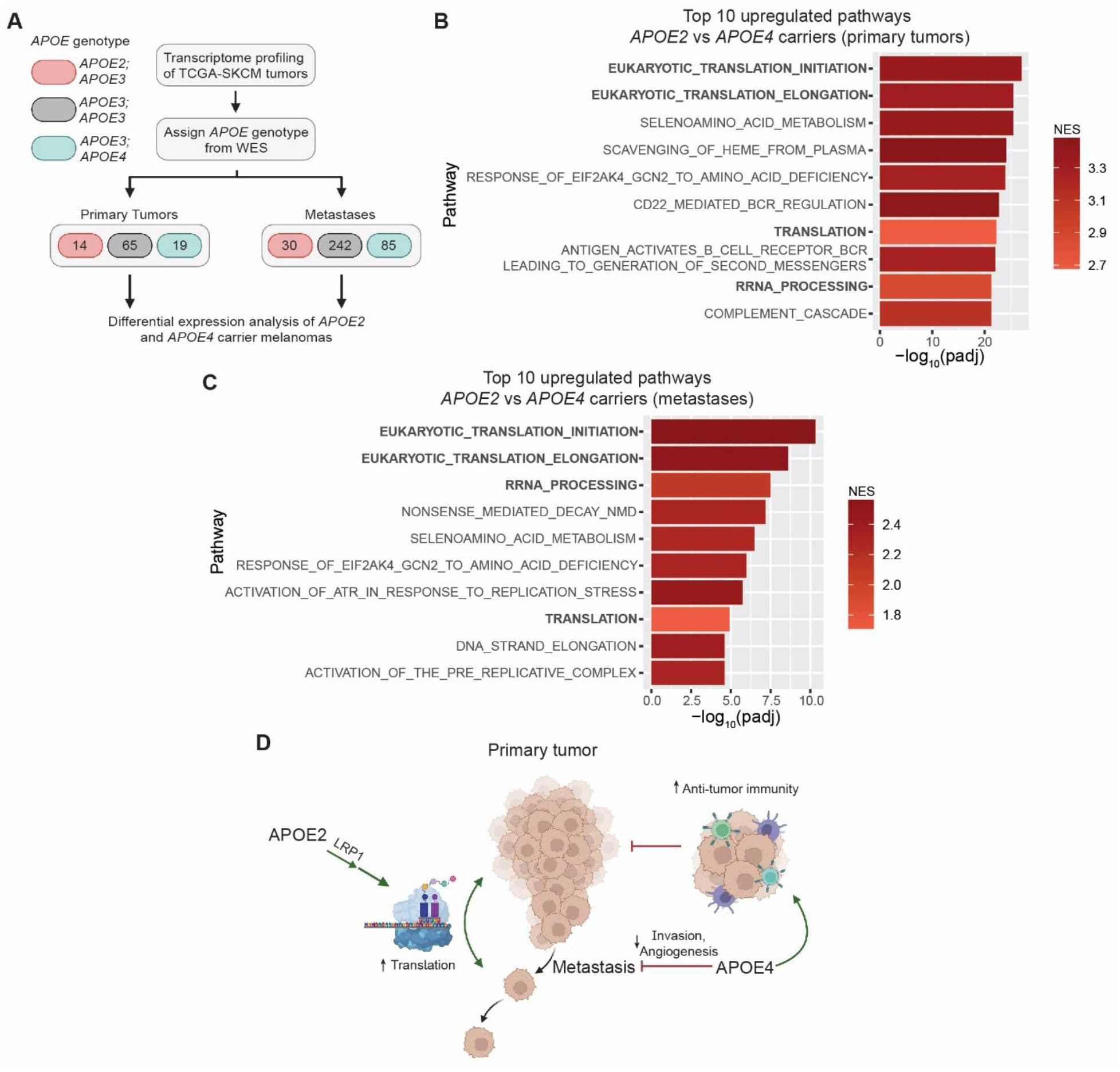
Protein translation pathways are upregulated in melanomas of *APOE2* carrier patients. **A**, Schematic depicting the workflow utilized to analyze transcriptomes of *APOE2* and *APOE4* carrier melanomas in the TCGA-SKCM cohort (WES, whole exome sequencing). **B**, Top ten pathways upregulated in primary tumors of *APOE2* carrier patients (n=14) relative to *APOE4* carrier patients (n=19) as determined by GSEA and ranked by adjusted p-value (NES, normalized enrichment score; padj, adjusted p-value). **C**, Top ten pathways upregulated in metastases of *APOE2* carrier patients (n=30) relative to *APOE4* carrier patients (n=85) as determined by GSEA and ranked by adjusted p-value. **D**, Model depicting our current understanding of the role of common *APOE* genetic variants in melanoma progression. The model depicts APOE4 acting as a suppressor of melanoma progression by enhancing anti-tumor immunity as well as repressing angiogenesis and invasion. In contrast, APOE2 is shown as a driver of melanoma progression through its stimulation of protein synthesis via the LRP1 receptor.

In sum, our findings uncover a new, surprising role of APOE and its genetic variants as differential modulators of mRNA translation in melanoma. We moreover provide evidence from a genetically initiated model that common germline variation in a human gene —*APOE*— differentially regulates tumor and metastatic progression. We propose a model in which APOE2 and APOE4 carry out opposing functions in melanoma, with APOE2 promoting growth and metastasis as well as protein translation via the LRP1 receptor and APOE4 repressing progression via its suppression of multiple cancer phenotypes including invasion, angiogenesis and immune evasion, as demonstrated in previous work (3) **(Fig. 4D)**.

## Discussion

Despite nearly 30 years of research following the discovery that *APOE* genotype impacts Alzheimer’s disease risk, the molecular mechanisms underlying this link remain elusive (7). This is a product of the pronounced complexity of APOE, which has diverse roles in numerous biological processes including lipid metabolism, immunity, mitochondrial function, and neuronal repair (7,20). It has also been unclear whether the *APOE* alleles represent a progressive gain or loss of function of the APOE protein, as the risk of APOE-influenced diseases other than Alzheimer’s does not always follow an *APOE4*>*APOE3*>*APOE2* order (7). Our study adds to an increasing body of evidence that the pleiotropic behavior of APOE also applies to melanoma (3,4) and supports the notion that *APOE2* can act as a gain-of-function allele in a disease context. We reveal that the APOE variants exert cell-intrinsic effects on melanoma that impact primary tumor growth and metastatic capacity in a *Braf*^*V600E*^-driven GEMM, a model that has provided numerous important insights into melanoma biology (21-25). Tumors in this melanoma GEMM progressed in an *APOE2*>*APOE3*>*APOE4* manner, consistent with melanoma patient survival outcomes. We expand the mechanism of this differential modulation beyond anti-tumor immunity, angiogenesis, and cellular invasion by identifying mRNA translation as an oncogenic process that is actively enhanced by APOE2.

Supporting the idea that our findings may have broader disease implications, our examination of previously published transcriptomic and proteomic Alzheimer’s datasets revealed that mRNA translation is a pathway that is upregulated in the brains of *APOE2* carriers relative to those of *APOE3* homozygotes and *APOE4* carriers (26,27). Neuronal protein synthesis has been well established as crucial for synaptic function and memory formation, and it is dysregulated in Alzheimer’s (28). Thus, enhancement of translation may contribute to the protective effect of APOE2 in Alzheimer’s and may partly explain the inverse impact of APOE variants in melanoma versus Alzheimer’s disease. Indeed, additional mechanistic connections between melanoma and Alzheimer’s disease are being increasingly uncovered (29).

We identified the APOE receptor LRP1 as necessary for the pro-tumorigenic and pro-metastatic activity of APOE2. Deletion of *Lrp1* abrogated differences in tumor growth, metastasis, and protein synthesis between *APOE2* and *APOE4* mice. LRP1 is a well-established regulator of intracellular signaling (30), thus making it suitable for mediating signals that could impact protein synthesis. Future studies are needed to elucidate the molecular mechanism downstream of this APOE2/LRP1 axis, as it may reveal new therapeutic targets for the treatment of melanoma and perhaps Alzheimer’s disease. Taken together, our work further highlights how germline genetic variation in the *APOE* gene impacts melanoma progression through disparate mechanisms. Although long speculated to exist, causal somatic mutational drivers of metastasis have not been identified despite extensive sequencing efforts, suggesting that alternative mechanisms may underlie propensity for metastatic progression (31). Our findings provide support for germline genetic variants as causal contributors to metastatic outcomes—revealing hereditary genetics as a predictor of cancer progression and providing new avenues for mechanistic and therapeutic studies.

## Materials and Methods

### Mice

Humanized *APOE2* (#1547, C57BL/6NTac), *APOE3* (#1548, C57BL/6), and *APOE4* (#1549, C57BL/6NTac) knock-in mice were obtained from Taconic Biosciences. *Braf* ^*V600E/+*^*;Pten*^*−/ −*^*;Tyr::CreER* (BPC) mice (RRID:IMSR_JAX:013590, C57BL/6J) were obtained from The Jackson Laboratory. *Lrp1*^*flox/flox*^ mice (C57BL/6J) were generously provided by David Hui (32). BPC mice were crossed with *APOE* knock-in mice to generate BPC/APOE2, BPC/APOE3, and BPC/APOE4 mice. BPC/APOE2 and BPC/APOE4 mice were crossed with *Lrp1*^*flox/flox*^ mice to generate BPC/APOE2/LRP1Δ and BPC/APOE4/LRP1Δ mice. Crosses were maintained on a C57BL/6J background.

### Mouse Genotyping

Genotyping of *Braf* ^*V600E/+*^*;Pten*^*−/ −*^*;Tyr::CreER* mice was performed as instructed by The Jackson Laboratory. Genotyping for *Lrp1*^*flox/flox*^ and discernment between mouse (200 bp) and human (∼600 bp) *APOE* was performed using standard PCR protocols. To distinguish between human *APOE* alleles, restriction fragment length polymorphism (RFLP) genotyping was performed (33). Briefly, a 244 bp portion of *APOE* was amplified using standard PCR protocols and digested simultaneously with AflIII (R0541) and HaeII (R0107) restriction enzymes (New England Biolabs) for at least two hours at 37°C. Allele-specific banding was visualized on a 4% agarose gel. The following PCR primers were utilized:

*Braf* ^*V600E/+*^*;Pten*^*−/ −*^*;Tyr::CreER genetic model*

Cre transgene forward: 5’ – GCG GTC TGG CAG TAA AAA CTA TC – 3’

Cre transgene reverse: 5’ – GTG AAA CAG CAT TGC TGT CAC TT – 3’

Cre internal control forward: 5’ – CAC GTG GGC TCC AGC ATT – 3’

Cre internal control reverse: 5’ – TCA CCA GTC ATT TCT GCC TTT G – 3’

Braf forward: 5’ – TGA GTA TTT TTG TGG CAA CTG C – 3’

Braf reverse: 5’ – CTC TGC TGG GAA AGC GGC – 3’

Pten forward: 5’ – CAA GCA CTC TGC GAA CTG AG – 3’

Pten reverse: 5’ — AAG TTT TTG AAG GCA AGA TGC — 3’

*Mouse versus human APOE*

Common forward: 5’ – TAC CGG CTC AAC TAG GAA CCA T – 3’

Mouse Apoe reverse: 5’ – TTT AAT CGT CCT CCA TCC CTG C – 3’

Human APOE reverse: 5’ – GTT CCA TCT CAG TCC CAG TCTC – 3’

*Human APOE allele RFLP*

Human APOE forward: 5’ – ACA GAA TTC GCC CCG GCC TGG TAC AC – 3’

Human APOE reverse: 5’ – TAA GCT TGG CAC GGC TGT CCA AGG A – 3’

*Lrp1*^*flox/flox*^

Lrp1 forward: 5’ – CAT ACC CTC TTC AAA CCC CTT CCT G – 3’

Lrp1 reverse: 5’ – GCA AGC TCT CCT GCT CAG ACC TGG A – 3’

### Cell Lines

HEK293T cells were obtained from the American Tissue Type Collection (ATCC). The B16F10 cell line transduced with a retroviral construct to express luciferase and GFP (34) and short hairpin RNA (shRNA) targeting murine *Apoe* (Millipore Sigma, TRCN0000011799; B16F10-TR-sh*Apoe*) was previously described (3,5). B16F10-TR-sh*Apoe* and HEK293T cells were cultured in DMEM (Gibco, 11995) supplemented with 10% fetal bovine serum (FBS) (D10F). All cells were maintained in an incubator at 37°C and 5% CO_2_ and regularly tested for *Mycoplasma* contamination with the Universal Mycoplasma Detection Kit (ATCC, 30-1012K).

### Generation of stable cell lines

APOE coding sequences from pCMV4-APOE2 (RRID:Addgene_87085), pCMV4-APOE3 (RRID:Addgene_87086), and pCMV4-APOE4 (RRID:Addgene_87087) plasmids were subcloned into the pBabe-hygro vector (RRID:Addgene_1765). Retrovirus was produced in HEK293T cells grown in 10 cm plates. Cells were transfected with retroviral Gag-pol (8 ug) and VSV-G (4 ug) packaging plasmids and pBabe vector (8 ug) using PEI Max transfection reagent (VWR, 75800-188). After 24 h, the medium was replaced with fresh DF10, and virus-containing supernatant was collected 48 and 72 h after transfection. The supernatant was filtered through a 0.45 µm filter, and viral supernatant was used with 8 µg/ml polybrene (Sigma, TR-1003-G) to transduce pre-plated B16F10-TR-sh*Apoe* cells for 8 hours. Following a second round of transduction, antibiotic selection was performed using 600 µg/ml Hygromycin B (Invitrogen, 10687010). Protein overexpression was validated by western blot.

### Generation of CRISPR Cell Lines

Single guide RNA sequences targeting murine *Lrp1* were obtained from the GeCKO v2 library (35) and cloned into the pSpCas(BB)-2A-Puro (PX459) V2.0 vector (RRID:Addgene_62988). B16F10-TR-sh*Apoe* cells were plated into a 6-well dish the day prior to transfection and transfected with 4 μg plasmid and TurboFect transfection reagent (Thermo Scientific, R0533) diluted in serum-free media, according to the manufacturer’s instructions. 24 hours after transfection, puromycin selection was initiated at 2 μg/mL concentration (Thermo Scientific, A1113803). Single clones were obtained by limiting dilution followed by Sanger sequencing of individual clones to confirm the presence of indels. Knockout clones were pooled to reconstitute heterogeneity, and LRP1 knockout was confirmed via western blot.

Guide RNA sequences:

sgCtrl: 5’ – GCGAGGTATTCGGCTCCGCG – 3’

sg*Lrp1*-1: 5’ – CCCGTTGCAGAGACGAGACA – 3’

sg*Lrp1*-2: 5’ – TTTGACGAGTGTTCCGTGTA – 3’

### Tail vein metastasis assay

6–8-week-old male *APOE2* and *APOE4* knock-in mice were injected via lateral tail vein with 100 µl of PBS containing 1 × 10^5^ B16F10-TR-sh*Apoe* cells. D-luciferin (GoldBio, 115144-35-9) was injected retro-orbitally, and bioluminescence was measured with an IVIS Lumina II (Caliper Life Sciences). Bioluminescence imaging was performed weekly, and signal was normalized to the signal obtained on day 0.

### Histology

Mice were perfused via intracardiac injection with PBS followed by 4% paraformaldehyde (PFA). The lungs were resected, incubated in 4% paraformaldehyde at 4°C overnight, and dehydrated in 70% ethanol at 4°C. Lungs were then embedded in paraffin, cut into 5 μm sections, and stained with hematoxylin and eosin (Histoserv, Inc.). Slides were digitally scanned with a PathScan Enabler (Meyer Instruments).

### Genetic tumor initiation

#### Topical induction

10 mg/ml of 4-hydroxytamoxifen (4-OHT; Sigma, H6278) was dissolved in acetone with gentle heating. 6–8-week-old female mice were shaved on the back, and 5 μl of 4-OHT was applied to back skin and allowed to air dry. Mice were observed twice weekly for tumor formation, defined as a raised, pigmented lesion at the site of tamoxifen application. Tumor volume was measured as described previously (21) by assessing length, width and height using a digital caliper (*V*=*l* x *w* x *h*), as tumors tended to grow cuboidal rather than spherical. For survival analyses, mice were euthanized according to the humane endpoints outlined in the IACUC protocol.

#### Perinatal induction

Two-day-old female neonates were tail snipped as described previously (36) and genotyped. 10 μl of 4-OHT diluted in DMSO (50 mg/mL) was applied with a small paintbrush to the back skin of neonates on postnatal days 3, 5, and 7. Mice were euthanized on postnatal day 35 or when moribund, whichever occurred earlier.

### Immunofluorescence of BPC tumor sections

Fresh tumors were excised, embedded in optimal cutting temperature (OCT) compound (Sakura Finetek, 4583), flash frozen in liquid nitrogen, and stored at -80°C. 20 µM tumor sections were obtained with a cryostat. Sections were fixed at -20°C with acetone/methanol and permeabilized with 0.1% Triton-X for 10 minutes at room temperature (RT). Blocking was performed for 30 minutes at RT with 5% goat serum in PBS with 0.1% Tween 20 (PBST). Sections were incubated at 4°C overnight with LRP1 primary antibody diluted in blocking solution (1:100; abcam 92544). Slides were washed with PBS and then incubated with Alexa Fluor 488 anti-rabbit secondary antibody diluted in PBST for 45 minutes (1:200; Invitrogen A11008). Slides were washed again with PBS, and nuclei were stained with 1 µg/mL of DAPI (Roche, 10236276001) followed by mounting with ProLong Gold Antifade Mountant (Invitrogen, P36930). Four independent fields per tumor section were imaged at random with a Nikon A1R MP confocal microscope with consistent instrument settings between samples. Sections stained with secondary antibody alone were used as negative controls.

### Quantification of BPC lung metastases

Lungs were fixed and dehydrated as described above. Lungs were then visualized with an OMAX trinocular microscope (W43C1-L08-TP). The number of pigmented lesions on the surface of each lung was quantified in a blinded manner under high magnification.

### Western Blot

Cells were lysed in ice cold RIPA buffer (G-Biosciences, 786-490) supplemented with protease inhibitor cocktail (Roche, 11836153001). Samples were denatured, separated by SDS-PAGE with 4-12% Bis-tris gels (Sigma), and transferred to low fluorescence PVDF membranes with the Trans-Blot Turbo Transfer System according to manufacturer’s instructions (Bio-Rad). Membranes were blocked for one hour with Intercept Blocking Buffer (LI-COR, 927-7000) and probed overnight at 4°C with the following primary antibodies diluted in blocking buffer containing 0.2% Tween-20: Puromycin (1:10000, Millipore #MABE343), APOE (1:1000, GeneTex #GTX100053), HSC70 (1:1000, Santa Cruz #sc-7298), LRP1 (1:50000, abcam #92544). Membranes were washed with PBST and incubated for an hour with IRDye 680RD goat anti-mouse (926-68070) or 800CW goat anti-rabbit (926-32211) secondary antibodies diluted in blocking buffer containing 0.2% Tween 20 and 0.1% SDS (LI-COR). Blots were imaged and analyzed with Image Studio Lite and Empiria Studio software (LI-COR).

### SUnSET Assay

#### In Vivo

BPC tumors were topically induced in mice as described above. 35 days after induction, mice were weighed and injected intraperitoneally with 40 nmol/g of puromycin. Mice were placed back in their cage for 30 minutes, after which mice were anesthetized with isoflurane and sacrificed by cervical dislocation. Tumors were dissected and rinsed with PBS to remove blood. ∼10mg of tumor was dissected from the center of tumors and homogenized with a Bead Ruptor Elite (Omni International) at 0°C in 200 μL of RIPA supplemented with protease inhibitor cocktail. To reduce viscosity, lysates were then treated with DNAse I (Norgen, 25710) according to manufacturer’s instructions. 40 μg of lysate was loaded and western blot was run and analyzed as described above. A mouse IgG2a-specific secondary antibody (1:5000; LI-COR, 926-32351) was used to eliminate background mouse IgG signal. Total protein was detected with the Revert 700 Total Protein Staining kit (LI-COR, 926-11010).

#### In Vitro

5×10^4^ B16F10-TR-sh*Apoe* cells stably expressing APOE2, APOE3, APOE4, or empty vector were plated in 8-well chamber slides (Nunc, 154941) the day before experiment. The next day, cells were serum starved for 6 hours in DMEM containing 0.2% FBS and then stimulated for 15 minutes with DF10. After stimulation, cells were pulsed with 10 μg/mL puromycin in DF10 for 30 minutes. As a positive control, one group of cells was treated with 100 μg/mL CHX for 10 minutes in DF10 prior to puromycin treatment. Cells were washed twice with PBS and then fixed with 4% PFA for 10 minutes. Cells were washed twice with PBS for 5 minutes each and then permeabilized with 0.5% Triton X-100 for 10 minutes. Two 5-minute PBS washes were performed, and cells were then incubated for 90 minutes at RT with 0.1% Triton X-100 containing Alexa Fluor 647-conjugated anti-puromycin antibody (1:5000; Millipore, MABE343-AF647). Cells were washed thrice with PBST for 5 minutes each. During the second wash, DAPI was added at a 1 μg/mL concentration. Slides were mounted with ProLong Gold and left to dry overnight. Four independent fields per condition were imaged at random with a Nikon A1R MP confocal microscope with consistent instrument settings between conditions. Mean fluorescence intensity was quantified with ImageJ.

### RNA extraction from BPC Tumors

Primary tumors were dissected 49 days after 4-OHT administration, flash frozen in liquid nitrogen, and stored at -80°C until RNA extraction. For RNA extraction, 10mg of tissue was dissected from the center of tumors on a ThermalTray (Corning, 432074) placed on dry ice. Tumor pieces were placed in homogenizer tubes containing ceramic beads along with lysis buffer from the Total RNA Purification Kit (Norgen, 37500) and RNAse inhibitors (10 μl/mL β-mercaptoethanol and 200 units/mL RNAsin Plus (Promega, N2615)), flash frozen in liquid nitrogen, and homogenized with a Bead Ruptor Elite at 0°C. RNA was then purified with the Total RNA Purification Kit with on-column DNAse treatment per manufacturer’s instructions.

### Digestion, purification, and RNA extraction of tail vein lung metastases

6–10-week-old female *APOE2* and *APOE3* knock-in mice were injected via lateral tail vein with 100 µl of PBS containing 1 × 10^5^ B16F10-TR-sh*Apoe* cells. Fifteen days after injection, mice were anesthetized via intraperitoneal injection of 2.5% Avertin in PBS (Sigma-Aldrich, T48402). Lungs were perfused with cold PBS intratracheally and via the left ventricle. Lungs were removed and placed in a 6-well plate on ice. Lungs were minced on ice with a scalpel and resuspended in 2mL of HBSS2+ (HBSS with calcium and magnesium (Gibco, 24020) supplemented with 2% FBS, 1 mM sodium pyruvate (Gibco, 11360), 25 mM HEPES (Gibco, 15630), 2 mg/mL collagenase IV (Worthington, LS004188), and 0.1 mg/ml DNAse I (Roche, 10104159001)) for 30 min at 37 °C on an orbital shaker at 80 rpm. 7 mL of HBSS2 − (HBSS without calcium and magnesium (Gibco, 14170), 2% FBS, 1 mM sodium pyruvate and 25 mM HEPES) was added to the digested tissue, which was homogenized with a serological pipette. The mixture was mashed through a 70 µm strainer and centrifuged at 500g for 5 min at 4°C. The supernatant was aspirated, and the pelleted cells were incubated in ACK lysis buffer (Lonza, 10-548E) for 3 minutes at RT. The mixture was neutralized with HBSS2 − and centrifuged. Cells were resuspended in 200uL of HBSS2 − and transferred to microcentrifuge tubes for flow sorting.

#### Flow Cytometry

Cells were centrifuged at 500g for 5 min at 4°C. Cells were then resuspended in 50 µl of flow buffer (1x PBS, 2% FBS, 10mM EDTA, 25mM HEPES) containing anti-CD16/CD32 antibody for Fc receptor blockade (Biolegend, 101320, 1:200) and incubated on ice for 5 minutes. For surface marker staining, 50 µL of flow buffer containing anti-CD45-BV785 antibody (Biolegend, 103149, 1:3000) was added, and samples were gently vortexed and incubated for another 20 minutes on ice under protection from light. Cells were then washed with flow buffer and resuspended in flow buffer containing TO-PRO-3 (Invitrogen, T3605). TO-PRO-3^low^/CD45^-^/GFP^+^ cells were sorted on a BD FACSAria II. Sorted tumor cells were centrifuged at 200g for 20 min at 4°C, and RNA was purified with the Total RNA Purification Kit with on-column DNAse treatment per manufacturer’s instructions. RNA was then concentrated with an RNA Clean & Concentrator Kit (Zymo Research, R1015).

### RNA-Seq

#### BPC Tumors

RNA integrity numbers (RIN) were measured with an Agilent Bioanalyzer 2100, with an average RIN of 8.15. TruSeq RNA Library Prep Kit v2, Set A (Illumina, RS-122-2001) was used to generate RNA-seq libraries according to manufacturer’s instructions. Libraries were quantified with an Agilent TapeStation and pooled at equimolar concentrations. Pooled libraries were sequenced with an Illumina NextSeq 500 (High Output, 75 SR). For analysis, FASTQ file quality was checked with FastQC (https://www.bioinformatics.babraham.ac.uk/projects/fastqc/). Kallisto v0.46.1 (37) was used to pseudoalign reads to the mm10 mouse transcriptome (version 101) downloaded from Ensembl. Quality data was aggregated with MultiQC (38). Counts were imported into R v4.1.3. with RStudio v2022.02.1 and tximport v1.18.0 (39). Differential expression analysis was performed with DESeq2 v1.28.1 (40) after prefiltering genes with less than 10 counts. Genes were annotated with AnnotationDbi v1.52.0 and the org.Mm.eg.db package. Genes were ranked based on Wald statistic, and GSEA was performed with the fgsea v1.20.0 (41). Mouse gene sets were downloaded from http://bioinf.wehi.edu.au/MSigDB/ based on MSigDB v7.1.

#### Tail vein lung metastases

RNA quality was assessed with an Agilent Bioanalyzer 2100, with an average RIN of 8.58. RNA-seq libraries were generated with a QuantSeq 3’ mRNA-Seq Library Prep Kit FWD (Lexogen) according to manufacturer’s instructions. Libraries were quantified, pooled, and sequenced as described above. FASTQ file quality was checked with FastQC. Adapter and poly A tail sequences were trimmed with bbduk v.38.31 (https://sourceforge.net/projects/bbmap/; options k=13, ktrim=r, forcetrimleft=11, useshortkmers=t, mink=5, qtrim=t, trimq=10, minlength=20). Reads were aligned to the GRCm38 mouse genome with STAR v.2.6.0 at default settings except for ‘outFilterMismatchNoverLmax’ set to 0.1 and ‘outFilterMultimapNmax’ set to 1. Mapped reads were counted with featureCounts (42). Differential expression analysis, gene annotation, and GSEA were performed as described above.

### TCGA Analysis

Harmonized raw counts of tumor transcriptomes from the TCGA-SKCM study were downloaded from the Genomic Data Commons API and imported into R with TCGAbiolinks v2.18.0 (43). *APOE* genotype information from whole exome sequencing was utilized as determined previously (3). Differential expression analysis was performed as described for BPC tumors, with tumor stage included as a covariate for primary tumor analysis to account for differences in tumor progression between genotypes. Genes were annotated with AnnotationDbi v1.52.0 and the org.Hs.eg.db package. The Reactome gene set was downloaded from MSigDB v7.1, and GSEA was performed as described above.

### Statistical Analysis

All data are expressed as mean ± SEM, unless indicated otherwise. Groups were compared using statistical tests for significance as described in the figure legends. A P value less than 0.05 was considered statistically significant. Statistical tests were performed with GraphPad Prism 9.

### Study Approval

All animal experiments were conducted in accordance with a protocol (#20010) approved by the Institutional Animal Care and Use Committee at The Rockefeller University.

## Data Availability

RNA-Seq data generated for this study have been deposited at the Gene Expression Omnibus under accession numbers GSE208718 and GSE209873. TCGA data is publicly available at the Genomic Data Commons: https://portal.gdc.cancer.gov/.

## Acknowledgements

We are grateful to members of the Tavazoie laboratory at Rockefeller University for feedback on the manuscript. We thank J. Posada for her pathology advice. We also thank the following Rockefeller University resource centers: V. Francis and other veterinary staff of the Comparative Bioscience Center, S. Mazel and staff of the Flow Cytometry Resource Center, C. Zhao and staff of the Genomics Resource Center, and staff of the Fisher Drug Discovery Resource Center. N.A. was supported by a Medical Scientist Training Program grant from the National Institute of General Medical Sciences of the National Institutes of Health under award number T32GM007739 to the Weill Cornell/Rockefeller/Sloan Kettering Tri-Institutional MD-PhD Program, an F30 Predoctoral Fellowship from the National Cancer Institute of the National Institutes of Health under award number F30CA257226, a medical student grant from the Melanoma Research Foundation, and the Shapiro-Silverberg Fund for the Advancement of Translational Research. S.F.T was supported by grants from the National Cancer Institute of the National Institutes of Health under award numbers R01CA184804 and U54CA261701 as well as the Black Family Metastasis Center. Schematics were created with BioRender.com.

## Author Contributions

**N. Adaku:** Conceptualization, data curation, formal analysis, funding acquisition, investigation, methodology, validation, visualization, writing – original draft, writing – review & editing

**B.N. Ostendorf:** Conceptualization, data curation, formal analysis, investigation, methodology, validation, writing – review & editing

**S.F. Tavazoie:** Conceptualization, supervision, funding acquisition, project administration, resources, writing – review & editing

