## Supplemental Figures for "Apolipoprotein E2 Promotes Melanoma Growth, Metastasis, and Protein Synthesis via the LRP1 Receptor"

### Supplementary Figures

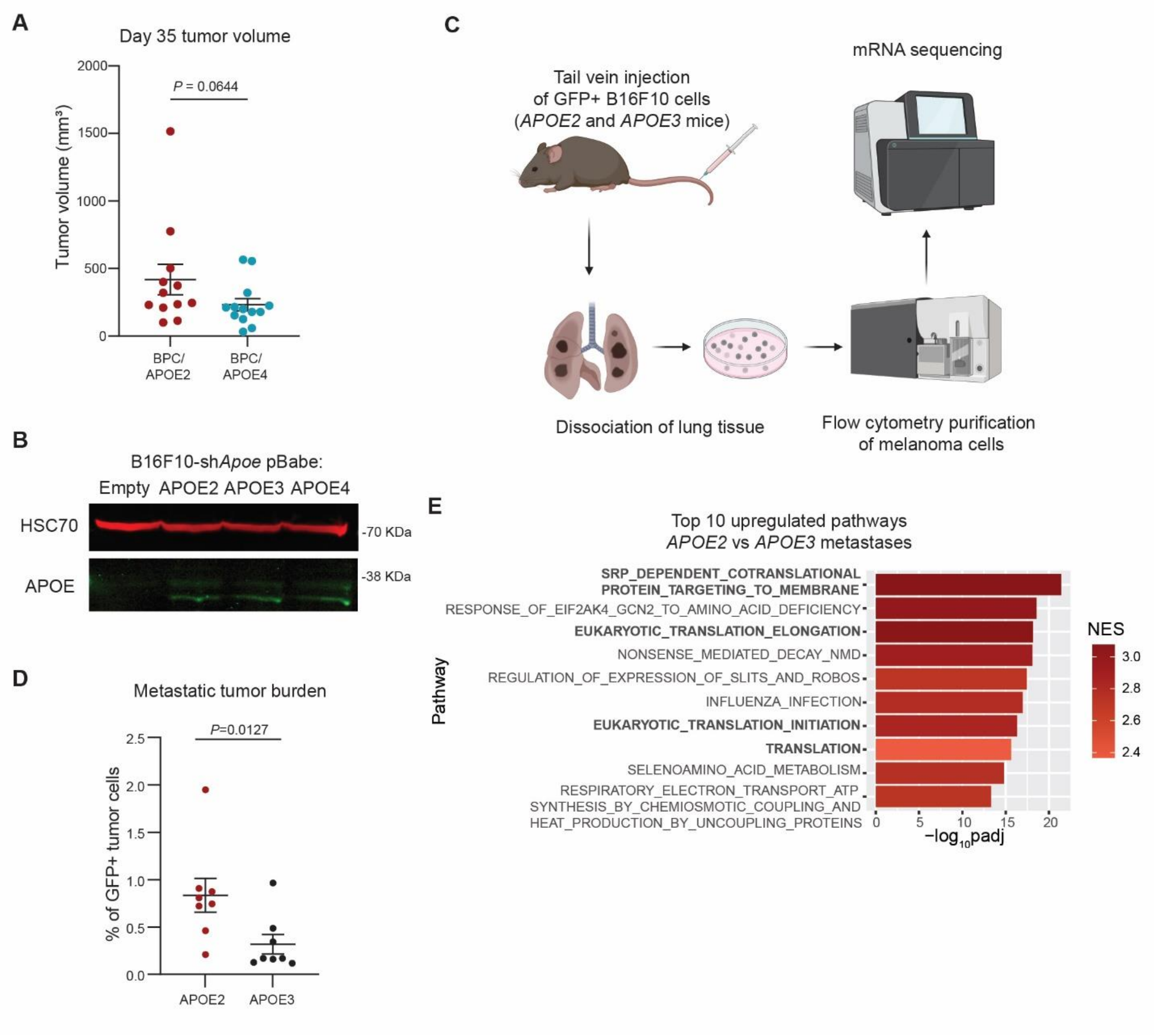


**Supplementary Figure 1. *APOE2* genotype promotes translation in melanoma**

**A,** Tumor volumes of BPC/APOE2 (n=12) and BPC/APOE4 (n=13) mice 35 days after 4-OHT administration. Unpaired t-test. **B,** Western blot of APOE expression in B16F10-sh*Apoe* cells transduced with pBabe Empty, APOE2, APOE3, or APOE4 retrovirus. HSC70 served as a loading control. **C,** Schematic depicting the workflow utilized to analyze transcriptomes of B16F10 lung metastases derived from *APOE2* and *APOE3* knock-in mice. **D**, Abundance of $GFP^+$ B16F10 tumor cells present in the total cell population derived from dissociated lung tissue of *APOE2* (n=8) and *APOE3* (n=8) knock-in mice. **E,** Top ten pathways upregulated in *APOE2* (n=8) lung metastases relative to *APOE3* (n=5) as determined by GSEA and ranked by adjusted p-value (NES, normalized enrichment score; padj, adjusted p-value)

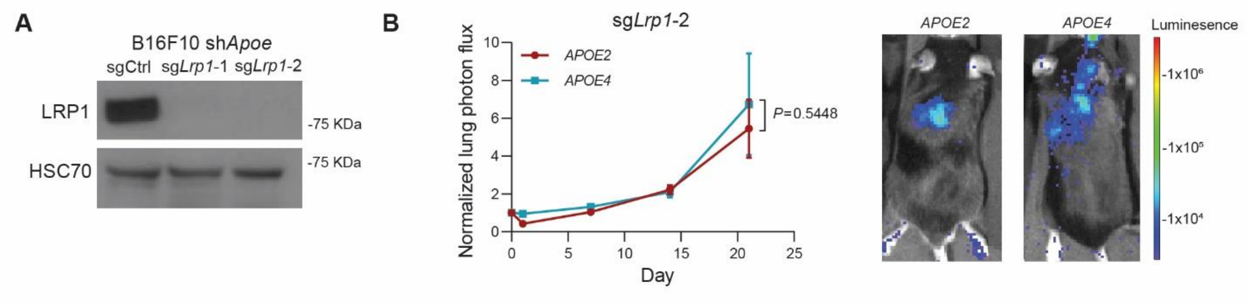


**Supplementary Figure 2. Tumoral *Lrp1* deletion abrogates *APOE* genotype-dependent differences in metastatic colonization**

**A,** Western blot of LRP1 expression in B16F10-TR-sh*Apoe* cells transfected with a non-targeting control CRISPR single guide RNA (sgRNA) or two independent *Lrp1*-targeting sgRNAs. HSC70 served as a loading control. **B,** Quantification of lung metastatic progression via bioluminescence imaging of B16F10-TR-sh*Apoe* sg*Lrp1*-2 cells injected via lateral tail vein into *APOE2* and *APOE4* mice. Representative bioluminescence images of mice at the day 21 endpoint. (n =8-10 mice per group; representative of two independent experiments; two-way ANOVA).

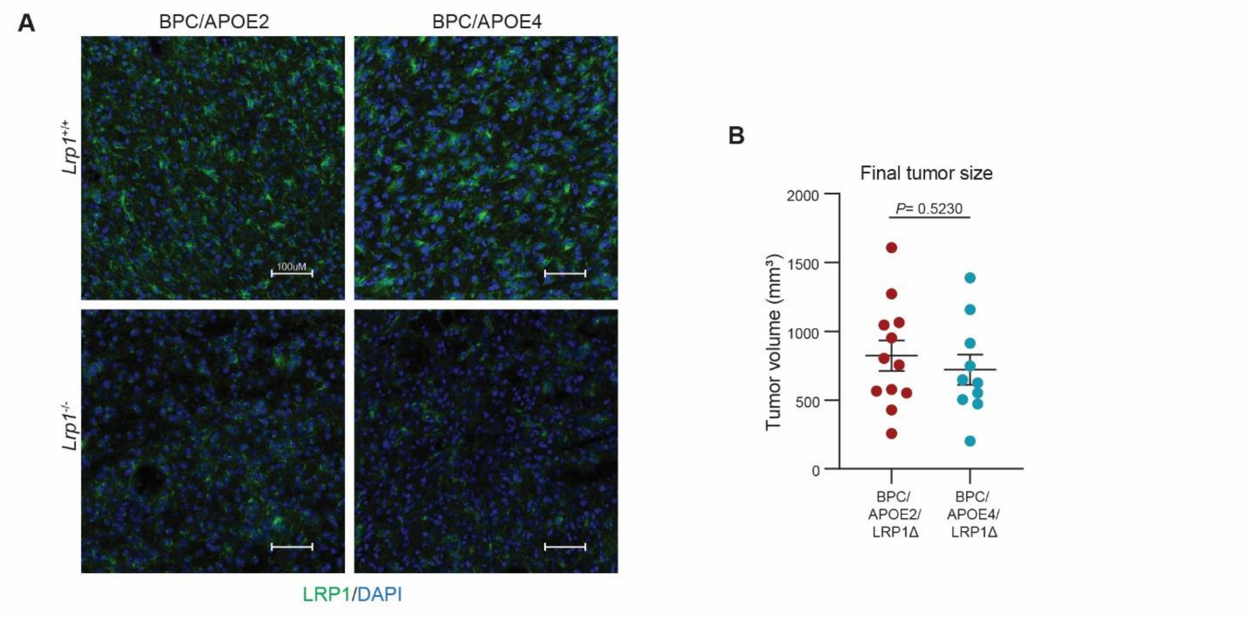


**Supplementary Figure 3. Tumoral LRP1 is a mediator of germline *APOE* variant differences in cancer progression and survival in a genetically initiated model of melanoma**

**A,** Representative immunofluorescence images of LRP1 expression and DAPI nuclear staining in BPC/APOE2, BPC/APOE4, BPC/APOE2/LRP1Δ, and BPC/APOE4/LRP1Δ primary tumors (scale bar = 100 μm). **B,** Final tumor volumes of BPC/APOE2/LRP1Δ (n=12) and BPC/APOE4/LRP1Δ (n=10) mice at the experimental endpoint 49 days after topical 4-OHT administration. Unpaired t-test.

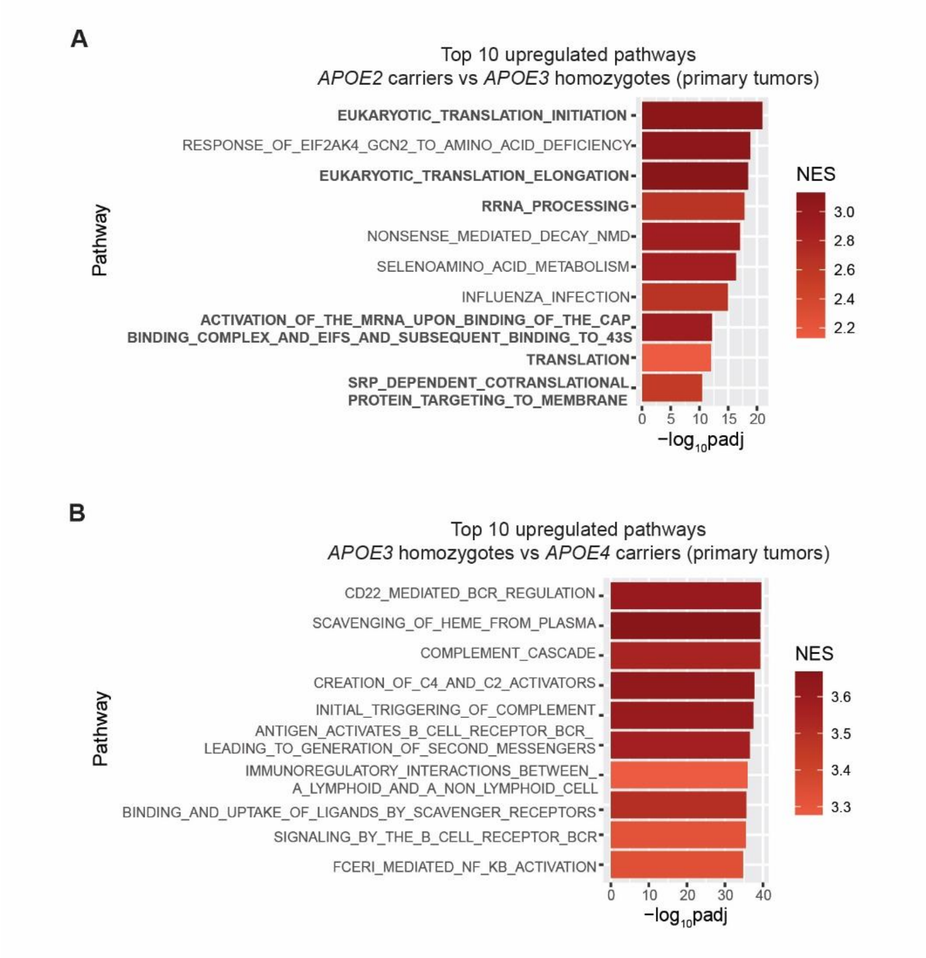


**Supplementary Figure 4. *APOE2* genotype associates with enrichment of mRNA translation gene expression pathways in human melanoma**

**A,** Top ten pathways upregulated in primary tumors of *APOE2* carrier patients (n=14) relative to *APOE3* homozygotes (n=65) as determined by GSEA and ranked by adjusted p-value (NES, normalized enrichment score; padj, adjusted p-value). **B,** Top ten pathways upregulated in primary tumors of *APOE3* homozygotes (n=65) relative to *APOE4* carrier patients (n=19) as determined by GSEA and ranked by adjusted p-value.
